# The Parkinson’s Disease related mutant VPS35 (D620N) amplifies the LRRK2 response to endolysosomal stress

**DOI:** 10.1101/2023.12.02.569621

**Authors:** Katy R. McCarron, Hannah Elcocks, Heather Mortiboys, Sylvie Urbé, Michael J. Clague

**Affiliations:** Biochemistry, Cell and Systems Biology, Institute of Systems, Molecular and Integrative Biology, University of Liverpool, Crown St., Liverpool, L69 3BX, UK; Sheffield Institute for Translational Neuroscience (SITraN), University of Sheffield, 385a Glossop Road, Sheffield, S10 2HQ, UK; Department of Molecular Microbiology & Immunology, Oregon Health & Science University, Portland, OR, USA

**Keywords:** LRRK2, VPS35, retromer, lysosomes, Parkinson’s Disease

## Abstract

The identification of multiple genes linked to Parkinson’s Disease invites the question as to how they may cooperate. We have generated isogenic cell lines that inducibly express either wild-type or a mutant form of the retromer component VPS35 (D620N), which has been linked to Parkinson’s Disease. This has enabled us to test proposed effects of this mutation in a setting where the relative expression reflects the physiological occurrence. We confirm that this mutation compromises VPS35 association with the WASH complex, but find no defect in WASH recruitment to endosomes, nor in the distribution of lysosomal receptors, cation-independent mannose-6-phosphate receptor and Sortilin. We show VPS35 (D620N) enhances the activity of the Parkinson’s associated kinase LRRK2 towards RAB12 under basal conditions. Furthermore, VPS35 (D620N) amplifies the LRRK2 response to endolysosomal stress resulting in enhanced phosphorylation of RABs 10 and 12. By comparing different types of endolysosomal stresses such as the ionophore nigericin and the membranolytic agent LLOMe, we are able to dissociate phospho-RAB accumulation from membrane rupture.

## Introduction

Parkinson’s disease (PD) is a complex neurodegenerative condition, for which variants of more than 20 genes have been linked to risk, onset and progression [1]. The major challenge in the field is to understand the cellular pathways linked to the pathology and how these genes exert their influence upon them [2]. It is particularly compelling when multiple PD genes can be linked to the same pathway. The first connection in this protein jigsaw was made through the genetic association of PINK1 and PRKN using *Drosophila* models [3–5]. The corresponding proteins were later shown to function in co-ordinating the clearance of damaged mitochondria, with PRKN being a direct substrate of the kinase PINK1 [6, 7].

Several PD genes have since been shown to converge on the endolysosomal pathway, but their specific connections remain to be established [8]. One of these is leucine-rich repeat kinase-2 (*LRRK2*) for which mutations leading to increased kinase activity are the most common cause of autosomal dominant PD [9]. Another one is VPS35, a component of the retromer complex, for which a specific heterozygous mutation (D620N) has been linked to PD [10–12]. A sub-set of members of the small GTPase RAB family serve as physiological substrates for LRRK2 (e.g. RAB8, RAB10, RAB12) and highly specific phospho-RAB antibodies have been introduced, which provide a proxy read-out for LRRK2 activity [13–15]. These tools enabled the discovery that damage to the endolysosomal membrane system, through membranolytic agents or infection, triggers the activation of LRRK2 and the subsequent recruitment of ESCRT-III components to repair the membrane [16–18]. Many suggestions have been made for the patho-physiological role of VPS35 (D620N) that use known retromer-dependent pathways as their inspiration. In addition, a persuasive link between VPS35 and LRRK2 has been established, in that VPS35 (D620N) results in hyperactivation of LRRK2 [19, 20].

Here, we have set out to design an isogenic cell system that allows inducible expression of VPS35 or VPS35 (D620N) at levels that are close to the endogenous protein. We have used these to systematically test for associated properties, previously claimed in the literature, many of which derive from over-expression studies. We confirm the VPS35-dependent hyper-activation of LRRK2 under basal conditions and moreover show that this remains extant under conditions of endosomal membrane stress.

## Results

### Generation of VPS35 isogenic cell models

We considered that some of the cell biological findings relating to the VPS35 mutation D620N would benefit from bench-marking in a systematic manner using an isogenic cell pair. As PD is induced by a heterozygous mutation, one physiologically correlated configuration is to express equal amounts of mutant protein and endogenous wild-type (WT) protein. To accomplish this, we adopted an RPE1 Flp-In cell line, which allows doxycycline induced expression of genes inserted at a unique integration site. This allowed us to generate and characterise isogenic cells, with induction of either wild-type HA-VPS35 or HA-VPS35 (D620N) to levels equivalent to the endogenous protein (Figure 1A-C). Both HA-tagged proteins co-localised with endogenous retromer protein (VPS26) on punctate structures, many of which are associated with EEA1-positive sorting endosomes (Figure 1D). They also showed equal distributions between cytosol and membrane fractions of mechanically homogenised cells suggesting that general membrane association of VPS35 is not compromised by the mutation (Figure 1E).

**Figure 1:**
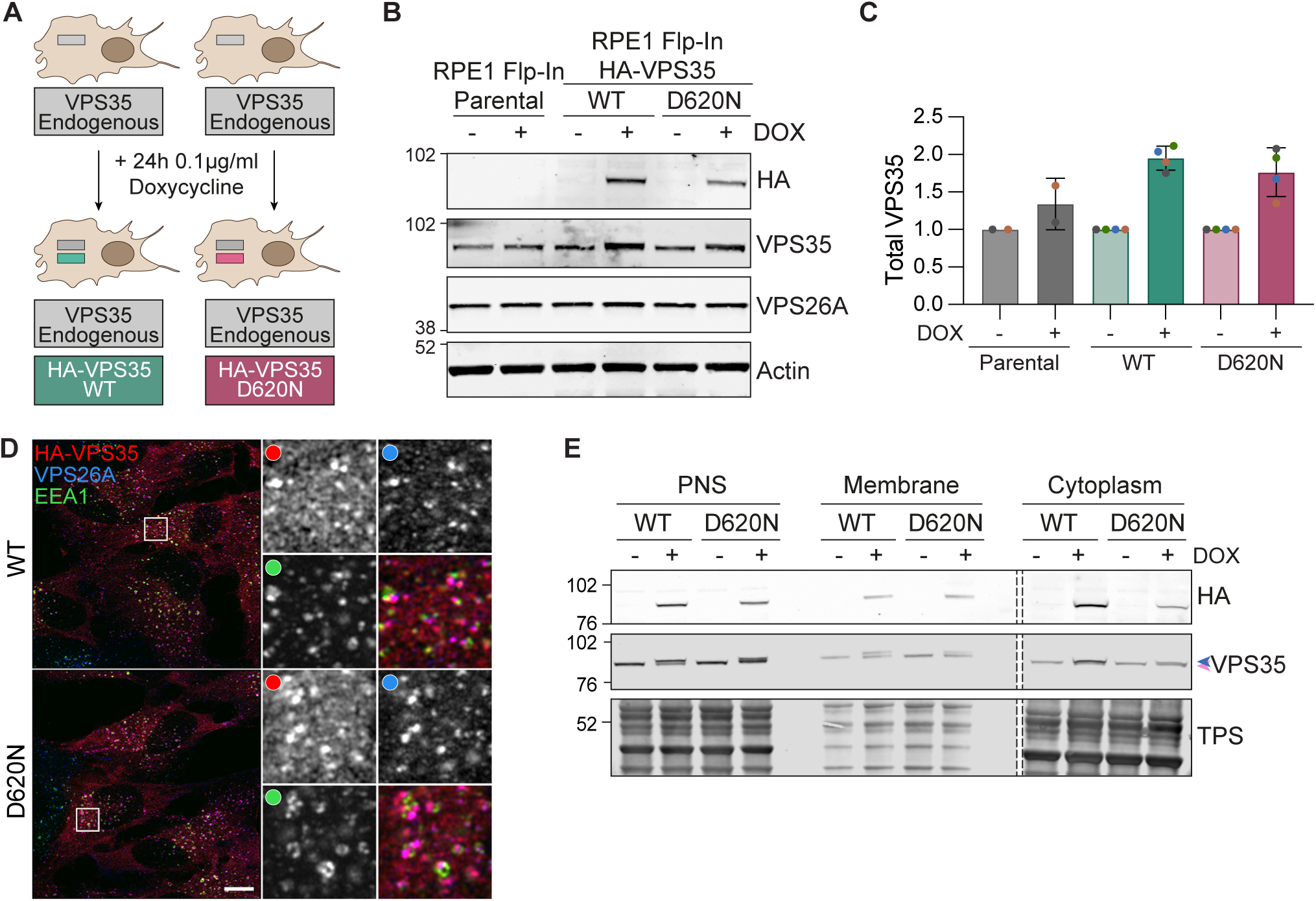
Characterisation of RPE1 Flp-In VPS35 isogenic cell lines. **A.** Schematic of doxycycline-inducible Flp-In system. **B.** Representative western blot of RPE1 Flp-In Parental, HA-VPS35 WT and HA-VPS35 (D620N) cells induced for HA-VPS35 expression with doxycycline (DOX) for 24 hours. **C.** Quantification of VPS35 expression normalised to actin then to “minus DOX control” for each cell line. n = 2-4 independent experiments. **D.** Representative Airyscan images of HA-VPS35 induced RPE1 Flp-In VPS35 WT and (D620N) cells stained for HA, VPS26 and EEA1. Scale bar 10 μm. **E.** Subcellular fractionation of RPE1 Flp-In VPS35 WT and (D620N) cells. Dashed line represents a loaded lane which has been cropped out. PNS, post nuclear supernatant; TPS, total protein stain.

### D620N does not compromise trafficking of lysosomal sorting receptors

VPS35 interacts with the WASH complex via interaction with FAM21 [21, 22]. Using a co-immunoprecipitation approach, we confirmed previous findings showing the VPS35 D620N mutation diminishes association with the WASH complex component, WASHC1, but not with the retromer component, VPS26 (Figure 2A,B) [20, 23, 24]. We could see no disruption to the recruitment of the WASH complex component FAM21 to VPS35 positive puncta, in line with the findings of McGough *et al.* (Figure 2 C-D) [23, 24]. We also found no change in the steady state distribution of lysosomal sorting receptors, cation-independent mannose-6-phosphate receptor (CIM6PR, otherwise known as IGFR2) and Sortilin, nor in the processing of a representative cargo, the lysosomal enzyme Cathepsin D (Figure 3). This is consistent with some previous reports [24, 25] and inconsistent with others (Table 1) [23, 26–28].

**Table 1.** comparison of results obtained with the VPS35 FlpIn cell model developed in this study with previous literature reports, relying on different contexts and configurations.

|  | [D620N] Mutant phenotype previously reported |  | RPE1 FlpIn VPS35 [D620N] model |
| --- | --- | --- | --- |
|  | Effect | Reference |  |
| LRRK2 activation | Enhanced | [19,20] | Enhanced |
| VPS35 association with WASH complex | Impaired | [23]<br>[24] | Impaired |
| Endosome localisation of WASH complex | Unaffected | [23] | Unaffected |
|  | Impaired | [24] |  |
| CIM6PR trafficking | Unaffected | [24, 25] | Unaffected |
|  | Impaired | [23, 26-28] |  |
| Sortilin trafficking | Unaffected | [25] | Unaffected |

**Figure 2:**
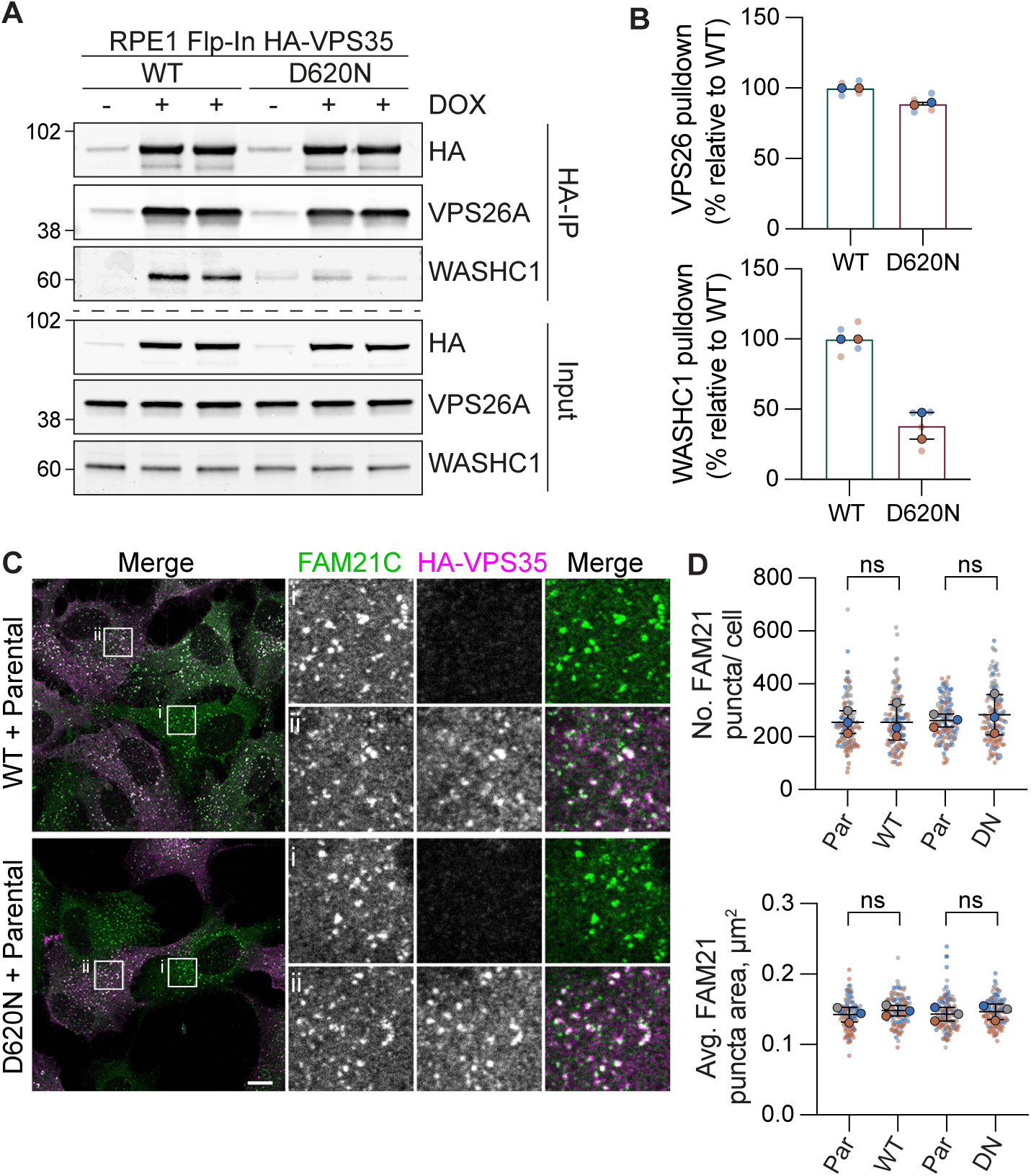
The VPS35 (D620N) mutation and WASH complex association. **A.** RPE1 Flp-In VPS35 WT and (D620N) cells were induced for HA-VPS35 expression with doxycycline and after 24 hours cell lysates were subject to immunoprecipitation (IP) using HA-antibody-coupled magnetic beads. IPs were probed alongside 3% of input, representative western blot is shown. **B.** Quantification of binding in the HA-IP relative to the mean of WT + DOX condition per experiment. n = 2 independent experiments with duplicate samples. Bars represent mean ± range. **C, D.** Representative images of RPE1 Flp-In VPS35 WT and (D620N) cells mixed with RPE1 Flp-In parental cells, induced with doxycycline for 72 hours then fixed and stained for FAM21 (C) and co-stained with an antibody against HA. Scale bar 10 *µ*m. (D) Quantification of FAM21 puncta, 33-50 cells counted per condition in each experiment. n = 3. One-way ANOVA with Tukey’s multiple comparisons test. ns, not significant.

**Figure 3:**
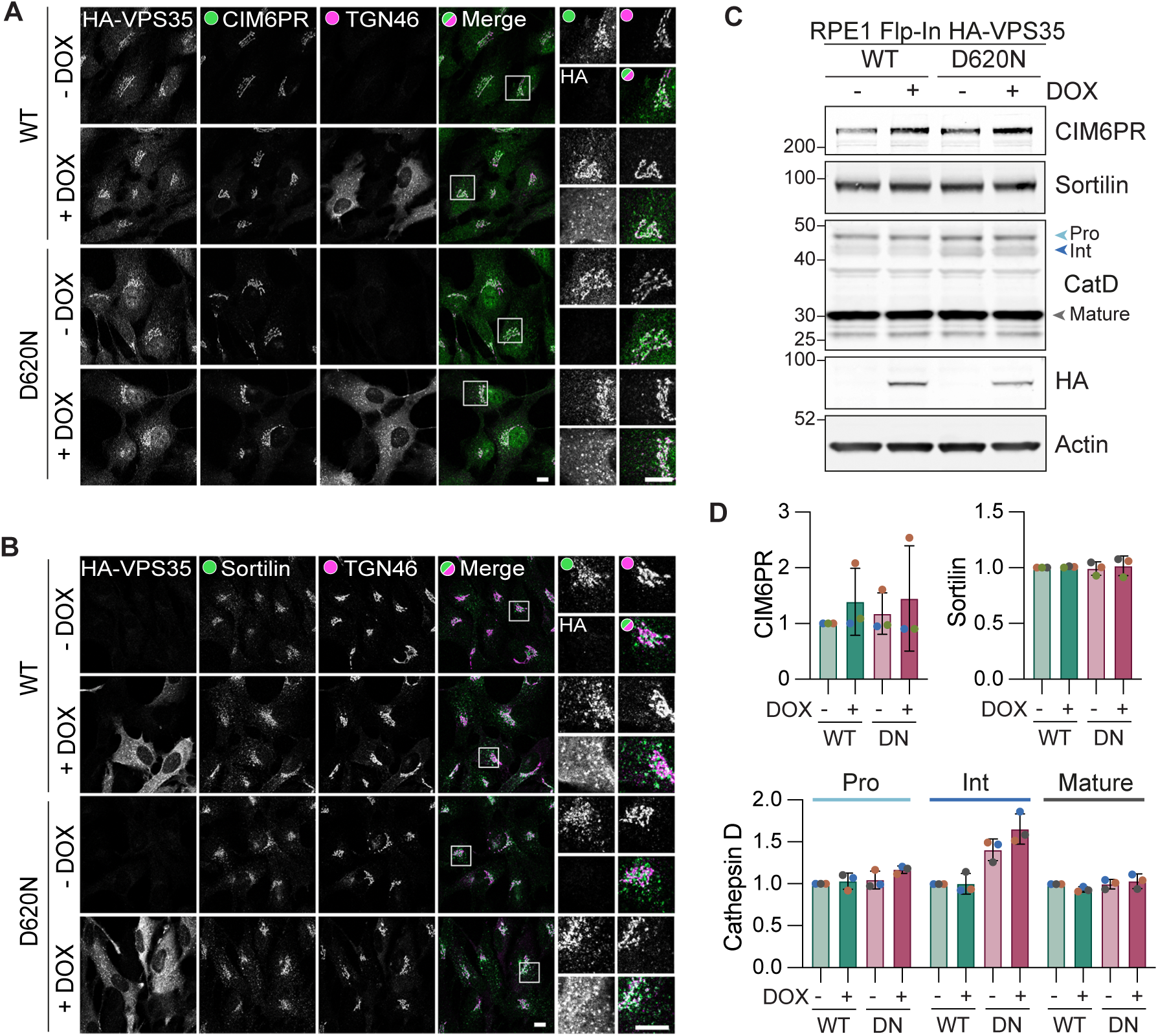
The VPS35 (D620N) mutation does not affect the trafficking of lysosomal receptors in RPE1 Flp-In VPS35 cells. **A, B.** Representative images of RPE1 Flp-In VPS35 WT and (D620N) cells induced with doxycycline (DOX) for 72 hours and then fixed and stained for CIM6PR (A) or Sortilin (B) and co-stained with TGN46 and HA. Scale bar 10*µ*m. **C.** Representative western blot of levels of Cathepsin D (CatD), CIM6PR and Sortilin in RPE1 Flp-In VPS35 WT and D620N (DN) cells. **D.** Quantification of C. Values normalised to WT minus DOX condition. n = 3.

### Interplay with LRRK2 and response to endolysosomal membrane damage

We next confirmed a previously reported synergy between the VPS35 D620N mutation and LRRK2 in our engineered cell system. Doxycycline-induced expression of VPS35 (D620N), but not an equivalent amount of wild type protein, lead to enhanced phosphorylation of RABs 10 and 12. This acts as a proxy read-out of LRRK2 activity and is accordingly sensitive to LRRK2 inhibitor MLi-2 (Figure 4A,B) [15].

**Figure 4:**
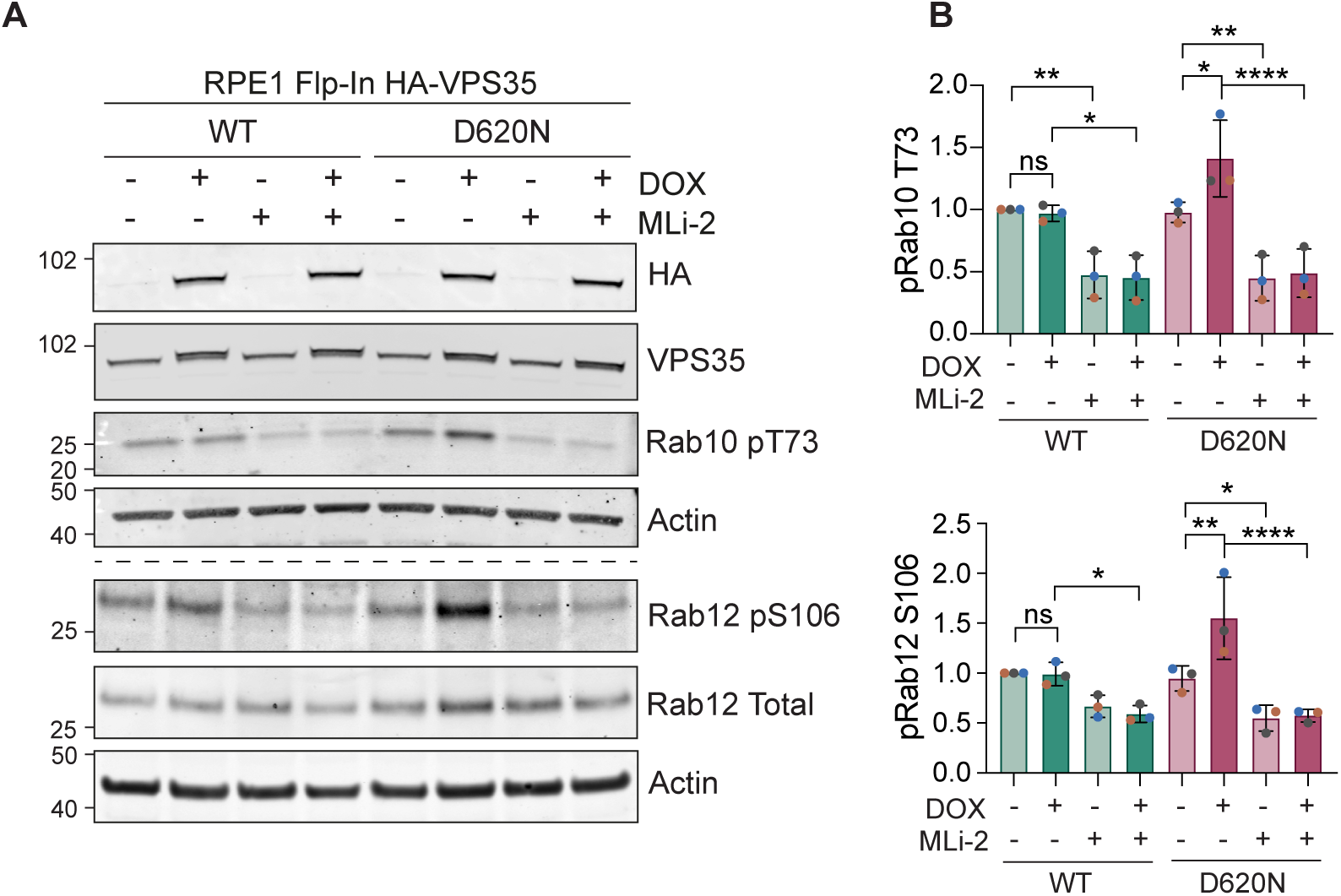
The VPS35 (D620N) mutation enhances basal Rab10 and Rab12 phosphorylation. **A.** Representative western blot of RPE1 Flp-In VPS35 WT and (D620N) cells induced for HA-VPS35 expression with doxycycline (DOX) and probed for phosphorylated and total Rab10 and Rab12. **B.** Quantification of phospho-Rab10 and 12 expression in A normalised to WT - DOX. Errors bars indicate mean ± SD. n = 3. One-way ANOVA with Tukey’s multiple comparisons test. P * < 0.05, P ** < 0.01, P *** < 0.001, P **** < 0.0001.

LRRK2 is activated by endolysosomal stress or damage, induced by invading pathogens or chemical manipulation [18, 29]. Common agents in use to accomplish this are chloroquine, nigericin and L-leucyl-leucine methyl ester (LLOMe). Nigericin is a K^+^/H^+^ exchanger which takes advantage of the high K^+^ gradient across the endosomal membrane and the osmotic inactivity of H^+^ ions to create an osmotic stress [30]. In contrast, LLOMe is converted into a membranolytic form by the action of cathepsins within the endolysosomal interior [31]. It has also been shown that endolysosomal osmotic stresses, including LLOMe and nigericin, can activate LC3-lipidation (LC3-II), mainly as part of a non-canonical autophagy pathway, referred to as conjugation of ATG8 to single membranes of the endolysosomal system (CASM) [32–36]. Here we show that the relative magnitudes of the response of two stress markers, pRAB12 and LC3-II in the parental cells, diverge according to the applied stress. pRAB12 is more elevated by nigericin, whilst LC3-II increases most strongly in response to LLOMe (Figure 5A,B). Furthermore, pRAB12 generation is contingent on the presence of endogenous VPS35, whilst LC3 lipidation is either insensitive or slightly enhanced (LLOMe) following VPS35 depletion (Figure 5A,Β). Nigericin-induced RAB phosphorylation is blocked by both inhibition of LRRK2 (MLi-2) or v-ATPase (concanamycin). Note that conconamycin alone does not stimulate RAB phosphorylation (Figure S1A,B). We also found that RAB phosphorylation and LC3-II generation are further enhanced by co-application of the PIKfyve inhibitor Apilimod together with nigericin (Figure S1A,B). This is consistent with a proposed mode of action, whereby it relieves a PtdIns3,5*P*_2_-dependent suppression of the Cl^-^/H^+^ antiporter CLC-7 [37]. Thus, osmotic stress will be further enhanced.

**Figure 5:**
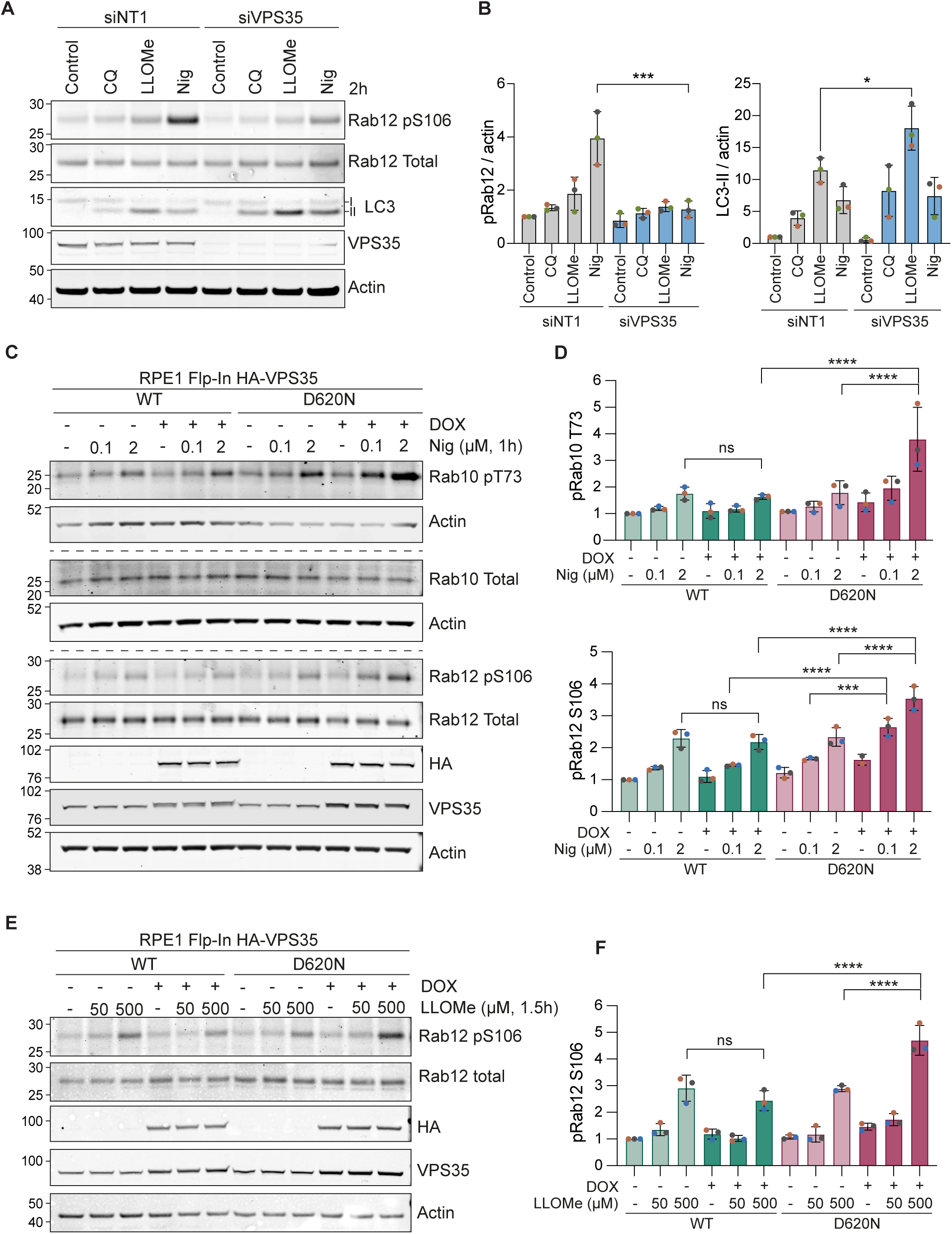
VPS35 is required for Rab phosphorylation in response to endolysosomal stress and is enhanced by the (D620N) mutation. **A.** Representative western blot of RPE1 Flp-In Parental cells treated with 40 nM of control (non-targeting #1; NT1) or VPS35 (pooled) targeting siRNA oligonucleotides for 72 hours and then treated with 2 *µ*M nigericin (Nig), 500 *µ*M LLOMe or vehicle (DMSO) for 2 hours prior to lysis. **B.** Quantification of A. Values normalised to actin then to the NT1 control sample. n = 3. Error bars represent mean ± SD. One-way ANOVA with Šídák’s multiple comparisons test performed on values normalised to sum of signal within a replicate then to actin. * P < 0.05, *** P < 0.001. **C.** Representative western blot of doxycycline (DOX)-induced RPE1 Flp-In VPS35 WT and (D620N) cells treated with 0.1 *µ*M or 2 *µ*M nigericin for 1 hour prior to lysis. **D.** Quantification of C. Values normalised to WT minus DOX control. Error bars indicate mean ± SD. n = 3. One-way ANOVA with Tukey’s multiple comparisons test. P *** < 0.001, P **** < 0.0001. **E.** Representative western blot of doxycycline-induced RPE1 Flp-In VPS35 WT and (D620N) cells treated with 50 *µ*M or 500 *µ*M LLOMe for 1.5 hours prior to lysis. **F.** Quantification of E. Values normalised to WT minus DOX control. Error bars indicate mean ± SD. One-way ANOVA with Tukey’s multiple comparisons test. P **** < 0.0001.

Although VPS35 (D620N) has previously been shown to stimulate basal LRRK2 activity this has not been examined in the context of endosomal stress [19]. We thus sought to examine if VPS35 (D620N) also amplified the LRRK2 response to endolysosomal damage. Whilst we saw small changes in the levels of basal RAB phosphorylation (RABs 10 and 12) following induction of VPS35 (D620N) alone, we found that this is highly amplified when endosomes are compromised by either nigericin or LLOMe (Figure 5C-F). All RAB phosphorylation induced by nigericin or LLOMe in the presence of VPS35 (D620N) is sensitive to LRRK2 inhibition by the specific inhibitor MLi-2 (Figure 6).

**Figure 6:**
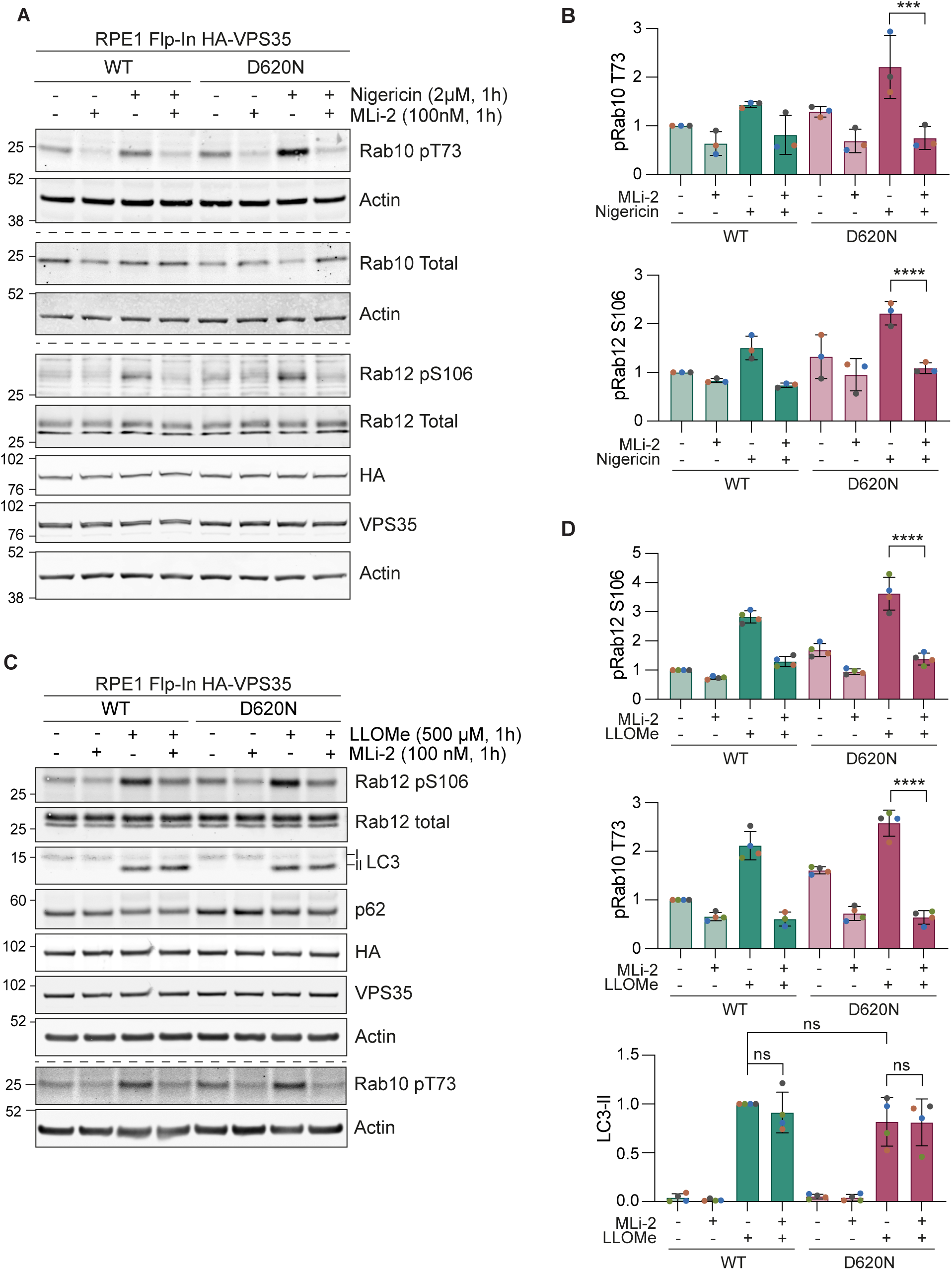
LRRK2 mediates (D620N) sensitised Rab phosphorylation. **A.** Representative western blot of doxycycline induced RPE1 Flp-In VPS35 WT and (D620N) cells co-treated with 2 *µ*M nigericin and 100 nM MLi-2 one hour prior to lysis. **B.** Quantification of A. Values normalised to the WT untreated sample. n = 3. Error bars indicate mean ± SD. One-way ANOVA with Tukey’s multiple comparisons test. P *** < 0.001, P **** < 0.0001. **C.** Representative western blot of induced RPE1 Flp-In VPS35 WT and (D620N) cells co-treated with 500 *µ*M LLOMe and 100 nM MLi-2 one hour prior to lysis. **D.** Quantification of C. Values normalised to the WT untreated sample. n = 4. Error bars indicate mean ± SD. One-way ANOVA with Tukey’s multiple comparisons test. P **** < 0.0001.

Endolysosomal membrane damage is known to lead to recruitment of components of the ESCRT-machinery that help repair breached membranes [38–41]. If this fails and endolysosomal membranes become leaky, they become accessible to Galectin 3, which binds to carbohydrate and co-ordinates a damage response [38]. This is evident following LLOMe but not nigericin treatment (Figure S1C,D) and is accompanied by recruitment of the ESCRT components CHMP2B and ALIX (Figure S1 E,F). We could not find any influence of VPS35 (D620N) upon recruitment of either CHMP2B or ALIX (Figure S2).

### VPS35 (D620N) limits clearance of LLOMe induced LC3-II

Following washout of previously applied LLOMe, endolysosomal membranes either reseal or undergo lysophagy as evidenced by a change in Gal3-mKeima emission that reflects acidification either due to membrane repair or lysophagy (Figure S1C- lower panel). After a 16 hour chase period, LC3-II positive puncta were greatly reduced. However, closer inspection of this residual signal reveals that there is a ∼2.5 fold increase of LC3-II in VPS35 (D620N) mutant cells compared with wild-type (Figure 7 A-C). Thus we show an effect of this VPS35 mutation on the recovery from membrane damage.

**Figure 7:**
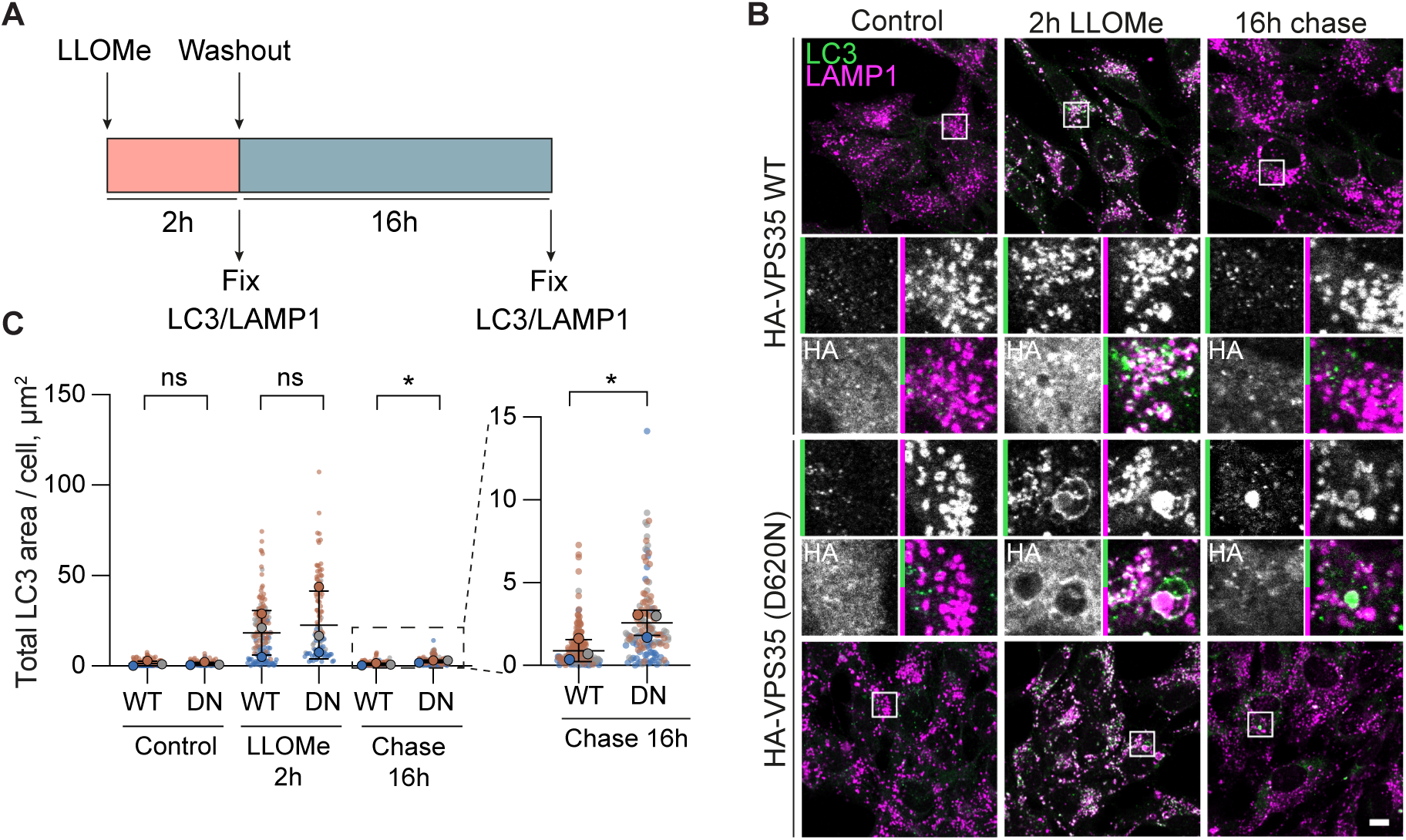
The (D620N) mutation impairs recovery from endolysosomal damage. **A.** Schematic of experimental set-up. **B.** Representative images of RPE1 Flp-In VPS35 WT and (D620N) (DN) cells induced with doxycycline for 24 hours and then treated with 500 *µ*M LLOMe or vehicle (DMSO) control for 2 hours. Cells were then either fixed or LLOMe was first chased out for 16 hours and then the cells were fixed and stained with the indicated antibodies. **C.** Quantification of the total area of LC3 puncta per cell. > 20 cells quantified per condition per repeat, *n* = 3. Error bars represent mean ± SD. Paired Student’s t-tests. * *P* < 0.05.

## Discussion

The cooperativity between VPS35 (D620N) and LRRK2 in the response to endosomal stress makes an important connection between PD-associated genes. Further links between PD and endolysosomal quality control are suggested by findings that lysosomal storage disorders can increase the risk of PD [42]. Most prominent amongst these is Gaucher’s disease for which 5-15% of patients will develop PD. Causative GBA1 mutations lead to loss of glucocerebrosidase activity and the accumulation of glucosylceramide and complex glycosphingolipids [43].

Here we introduce a new isogenic cell model for the study of a mutation in VPS35 (D620N), which reflects the heterozygous state occurring in PD. We have assessed previous findings in this system and our results are summarised in Table 1. The value of our data lies in testing these parameters in a unique setting that reflects the physiological expression levels. Where we fail to substantiate past claims, it could reflect different expression levels or simply biological context e.g. epithelial cells versus neurons. However, we can recapitulate two robust findings associated with the VPS35 (D620N) mutation i.e. diminished interaction between retromer and the WASH complex and activation of LRRK2 [19, 20, 23]. This second facet is the main focus of our study, as it reflects a direct connection between two pieces of the Parkinson’s Disease jigsaw.

We find that the expression of VPS35 (D620N), at physiological levels, leads to enhancement of LRRK2 activity in response to endolysosomal stress inducing agents. This study is the first to show that the LRRK2 response to endolysosomal stress is enhanced by VPS35 (D620N), in addition to the previously reported role of this mutant in promoting basal activity in mouse embryonic fibroblasts [19]. Thus the VPS35 mutation alone recapitulates key signatures of activating LRRK2 mutations found in PD.

LRRK2 activity allows for engagement of a membrane repair pathway, which may protect cells from toxic entities and stress [18]. By applying different endolysosomal stresses we have been able to dissociate some of the signature sequelae. The osmotic stress generated by nigericin alone or in combination with apilomod gives the strongest proxy read-out of LRRK2 activity, provided by pRAB10 and pRAB12. In contrast, the direct induction of membrane damage by LLOMe, perhaps the most widely used inducer of LRRK2 activity, gives a weaker pRAB response, but is accompanied by LC3 lipidation and the recruitment of ESCRT proteins for membrane repair. This initial LC3/ESCRT response to membranolytic damage is not influenced by VPS35 (D620N) expression. However, we do find an influence of VPS35 (D620N) in the recovery from LLOMe induced endolysosomal membrane damage, which merits future detailed exploration.

Our data indicate that membrane rupture *per se* is not required for LRRK2 activation. Likewise, all of the drugs we have used will dissipate the pH gradient, so that alone cannot account for our findings. This is directly supported by our finding that the v-ATPase inhibitor concanamycin has a minimal effect upon RAB phosphorylation. We propose that instead LRRK2 activation may principally be responding to increased membrane surface tension associated with osmotic stress or other factors. As RAB proteins control multiple facets of membrane trafficking, they may be marshalled by LRRK2 to alleviate this stress; for example, by increasing the limiting membrane area or by remodelling their composition to regulate the flux of osmolytes [17].

The present work highlights the importance of understanding the endolysosomal membrane properties leading to LRRK2 activation and any point of convergence. Comprehending the interplay between PD associated genes is the major challenge to generating a holistic framework for the molecular mechanisms of this disease. The Parkinson’s linked lysosomal ATPase, ATP13A2, has recently been shown to function as a H^+^/K^+^ and polyamine transporter linked to Parkinson’s disease [44–46]. It will be interesting to see how these regulate the turgor pressure within endolysosomal compartments and to explore the interplay with LRRK2 activity.

## Materials and Methods

### Antibodies and reagents

Antibodies and other reagents are as follows: anti-actin (1:10,000 WB, Proteintech; 66009-1-IG), anti-actin (1:2,000 WB, Proteintech; 20536-1-AP), anti-actin (1:2,000 WB, Sigma-Aldrich; A2066), anti-ALIX (1:1,000 WB, 1:500 IF, Santa Cruz Biotechnology; sc-53540), anti-Cathepsin D (1:2,000 WB, Calbiochem; 219361), anti-CD63 (1:500 IF, Developmental Studies Hybridoma Bank H5C6), anti-CHMP2B (1:500 IF, abcam ab157208), anti-CIM6PR (1:1,000 WB, 1:500 IF, gift from Paul Luzio), anti-EEA1 (1:500 IF, BD Biosciences; 610456), anti-FAM21C (1:1,000 IF, Merck; ABT79), anti-HA (1:1000 WB, Covance; MMS-101P), anti-HA (1:400 IF, Novus Biotechne; NB600-362), anti-HA (1:500 IF, Roche; 11867423001), anti-LAMP1 (1:1,000 WB, 1:500 IF, Cell Signalling Technology; 9091), anti-LAMP1 (1:200 IF, Developmental Studies Hybridoma Bank H4A3), anti-LC3 (1:200 WB, 1:200 IF, Nanotools; 5F10), anti-p62 (1:1,000 WB, BD Biosciences; 610833), anti-phosphoRab10 T73 (1 *µ*g/ml WB, abcam; ab230261), anti-phosphoRab12 S106 (1:1,000 WB, abcam ab256487), anti-Rab10 total (1:1000 WB, Cell Signalling Technology; 8127), anti-Rab12 total (1 *µ*g/ml WB, MRC PPU SA227), anti-Sortilin (1:1,000 WB, 1:1,000 IF, abcam; ab16640), anti-TGN46 (1:500 IF, Biorad; AHP500), anti-VPS26A IF, WB (1:1,000 WB, 1:800 IF, abcam; ab23892), anti-VPS35 (1:1000 WB, abcam; ab10099), anti-WASHC1 (1:2,000 WB, Sigma-Aldrich HPA002689), apilimod dimesylate (Tocris Bioscience; 7283), blasticidin S (Invitrogen; R21001), chloroquine diphosphate salt (Sigma-Aldrich; C6628), concanamycin A (Sigma-Aldrich; C9705), doxycycline hyclate (Sigma-Aldrich; R25005), G-418 solution (Roche; 4727878001), L-Leucyl-L-Leucine methyl ester (LLOMe) hydrochloride (Cayman Chemical; CAY16008), MLi-2 (Tocris Bioscience; 5756), nigericin sodium salt (Sigma-Aldrich; N7143), puromycin dihydrochloride (Sigma-Aldrich; P7255), Zeocin (Invitrogen; R25005).

### Cell culture, treatments and transfections

hTERT-RPE1-Flp-In-Parental (hTERT-RPE1-FRT-TREX, kind gift from Jon Pines, ICR, London) cells were cultured in Dulbecco’s modified Eagle medium DMEM/F12 with GlutaMAX (Gibco; 31331093) supplemented with 10% FBS (Gibco; 10270106) and 1% non-essential amino acids (Gibco; 11150035). hTERT-RPE1-Flp-In-HA-VPS35 cells were maintained as above in the presence of 200 *µ*g/ml G418 (Sigma; 4727878001) and 5 *µ*g/ml blasticidin (Thermo Fisher Scientific; R21001). Lenti-X HEK293T cells were cultured in Dulbecco’s modified Eagle medium supplemented with 10% FBS. Cell lines were tested regularly for mycoplasma. Apilimod, concanamycin A, L-Leucyl-L-Leucine methyl ester (LLOMe) and MLi-2 were solubilised in sterile DMSO. Chloroquine was solubilised in filtered sterile water. Nigericin was solubilised in absolute ethanol. Treatments were added at a dilution of 0.1% to cells. For siRNA-mediated knockdowns, cells were transfected with 40 nM siRNA using Lipofectamine RNAiMAX (Invitrogen; 13778075) in serum-free DMEM/F12 according to manufacturer’s protocol. Media was exchanged after 6 hours for fully supplemented DMEM/F12 and cells were harvested 72 hours after transfection. ON-TARGETplus Human VPS35 siRNA-SMARTpool (L-010894-00) and ON-TARGETplus non-targeting control #1 (NT1) siRNA (D-001810-01) were purchased from Horizon Discovery/Dharmacon.

### Generation of HA-VPS35 Flp-In cell lines

The expression plasmids pCDNA5 FRT/TO-NeoR-HA-VPS35 WT and [D620N] were generated by subcloning either pcDNA5D-FRT/TO-HA-VPS35-WT (MRC PPU; DU26467) or pcDNA5D-FRT/TO-HA-VPS35-D620N (MRC PPU; DU26878) into the vector pcDNA5/FRT/TO-neo (Addgene; 41000). Prior to transfection, hTERT-RPE1-Flp-In-Parental were maintained in 10 *µ*g/ml blasticidin and 100 *µ*g/ml zeocin. To generate stable cell lines, hTERT-RPE1-Flp-In-Parental cells were co-transfected with a 10:1 ratio of pOG44 and either pCDNA5 FRT/TO-NeoR-HA-VPS35-WT or -[D620N] using Genejuice (Merck Millipore; 70967), according to the manufacturer’s protocol. Clones were selected under 400 *µ*g/ml G418/geneticin (neomycin) and 10 *µ*g/ml blasticidin. Individual clones were amplified and validated by western blotting and immunofluorescence. VPS35 expression in RPE1-Flp-In cell lines was induced by doxycycline (0.1 *µ*g/ml) for 24 hours unless otherwise stated.

### Generation of lysophagy reporter RPE1 Flp-In HA-VPS35 mKeima-Gal3 cell lines

To produce lentivirus, 450,000 Lenti-X HEK293T (Clontech) cells were seeded per well of a 6-well plate and co-transfected with 2.8 *µ*g pHAGE-mKeima-LGALS3 (Addgene; 175780), 2.3 *µ*g pPAX2 (Addgene; 12260) and 0.85 *µ*g pMD2G (Addgene; 122259) using Lipofectamine 2000 (Invitrogen; 1668027), according to the manufacturer’s protocol. After 48 hours, the virus-containing media was harvested. To determine the viral titre and generate stably expressing cells, RPE1 Flp-In VPS35 WT and (D620N) cells were transduced with mKeima-LGALS3 (Gal3) using polybrene (8 *µ*g/ml) and after 24 hours, subjected to puromycin selection (1 *µ*g/ml). The mixed pools of puromycin-resistant cells were used for experiments.

### Preparation of cell lysates and Western blot analysis

Cultured cells were lysed in RIPA (150 mM NaCl, 1% sodium deoxycholate, 0.1% SDS, 1% Triton X-100, 10 mM Tris-HCl pH7.5) lysis buffer routinely supplemented with mammalian protease inhibitor cocktail (MPI; Sigma-Aldrich; P8340) and PhosSTOP (Roche; 4906837001). Proteins were resolved using SDS-PAGE (Invitrogen NuPage 4-12%), transferred onto nitrocellulose membrane (Amersham; 10600001 or 10600002), stained with Ponceau S staining solution (Sigma-Aldrich, P7170), blocked for 1 hour in 5% milk (Marvel) in TBS supplemented with 1% Tween-20 (TBST), then probed with primary antibodies. Primary antibodies were incubated in 5% milk /TBST, apart from phospho-antibodies, which were incubated in 5% BSA/TBST. Visualisation and quantification of Western blots was performed using IRdye 800CW and 680LT coupled secondary antibodies and an Odyssey infrared imaging system (LI-COR Biosciences). For western blot quantification, raw signal values were obtained using ImageStudio Lite (LI-COR) software following background subtraction. For measurement of statistical significance, raw values for each condition were normalised to the sum of the quantified raw values from each individual blot.

### Subcellular fractionation

RPE1 cells were washed twice with ice-cold PBS and then collected by scraping in PBS and pelleting at 1,000 *g* for 2 min. Cell pellets were washed with HIM buffer (200 mM mannitol, 70 mM sucrose, 1 mM EGTA, 10mM HEPES-NaOH pH7.4) then resuspended in HIM buffer supplemented with MPI and PhosSTOP. Cells were mechanically disrupted by shearing through a 23G needle. The lysed cells were centrifuged at 600 *g* for 10 min at 4 °C, and the post-nuclear supernatants (PNS) transferred. The PNS was separated into cytosolic and membrane fractions by centrifugation at 100,000 *g* for 30 min at 4 °C. The membrane pellet was resuspended in HIM buffer supplemented with MPI and PhosSTOP. Sample concentrations were normalised following BCA protein assay and analysed by western blotting. Total protein was visualised using Revert 700 Total Protein Stain (LI-COR; 926-11021).

### Immunofluorescence and live-cell imaging

Generally, cells were fixed and permeabilised with ice cold methanol for 5 minutes. For some experiments (Figures 1D, Fig 2C and Fig S1F), cells were fixed with 4% paraformaldehyde/PBS for 15 min, quenched with 50 mM ammonium chloride/PBS for 10 min, and permeabilised with 0.2% Triton X-100/PBS for 4 min. Coverslips were incubated at room temperature for 30 min in 10% goat serum (Sigma-Aldrich; G6767)/PBS or 3% BSA (41-10-410; First Link)/PBS blocking solution, followed by a 1 hour incubation with primary antibodies and 20 min with AlexaFluor-conjugated secondary antibodies (Invitrogen) in 5% goat serum or 3% BSA in PBS, before mounting in Mowiol with DAPI (Invitrogen; D1306). Images were acquired with a Zeiss LSM800 with Airyscan (63x NA 1.4 oil) or Zeiss LSM900 with Airyscan 2 (63x NA 1.4 oil) confocal microscope. For live-cell imaging of mKeima-Gal3 (lyso-Keima), cells were seeded into ibidi 8 well chamber *µ*-slides (ibidi; 80826) three days prior to imaging with a 3i Marianas spinning-disk confocal microscope (63x oil objective, NA 1.4, Photometrics Evolve EMCCD camera, acquisition software Slide Book 3i v3.0). Images were acquired sequentially (445 nm excitation, 617/73 nm emission; 561 nm excitation, 617/73 nm emission).

Image analysis was performed using Fiji (v 2.14.0). Cells were manually traced using the Freehand selection tool to generate regions of interest. Every field of view for each condition in an experiment was thresholded according to a consistent pipeline and the ‘Analyse particles’ tool was used to identify puncta and measure their average size, number and fluorescence intensity per cell. Images were processed using OMERO.web (v 5.5) and Adobe Photoshop (v 25.0.0) software.

### Statistical analysis

Bars indicate mean and standard deviation (SD) or range, as indicated. Individual data points are colour-coded according to each independent experiment. Transparent circles with no outline represent values for each sample in an experiment, opaque circles with or without black outlines correspond to mean values for each experiment. Statistical significance was determined with an unpaired t-test (Fig 7C) or one-way ANOVA with either Tukey’s (Figures 2D, 4B, 5D, 5F, 6B, 6D, S2B, S2D) or Šídák’s (Fig 5B) multiple comparisons tests using Graph Pad Prism 10. P-values are represented as *P < 0.05, **P < 0.01, ***P < 0.001 and ****P < 0.0001.

## Acknowledgements

KRM is supported by a studentship from the Medical Research Council (MRC, MR/ N013840/1) Discovery Medicine North (DiMeN) Doctoral Training Partnership and has received additional funding from the Professor John Glover Memorial Award. HE was supported by a studentship from the Medical Research Council (MRC) Discovery Medicine North (DiMeN) Doctoral Training Partnership and by a Parkinson’s UK Grant (G-1902). MC is a Royal Society Industry Fellow, INF\R2\212031.

## Supplementary Figure Legends

**Figure S1:**
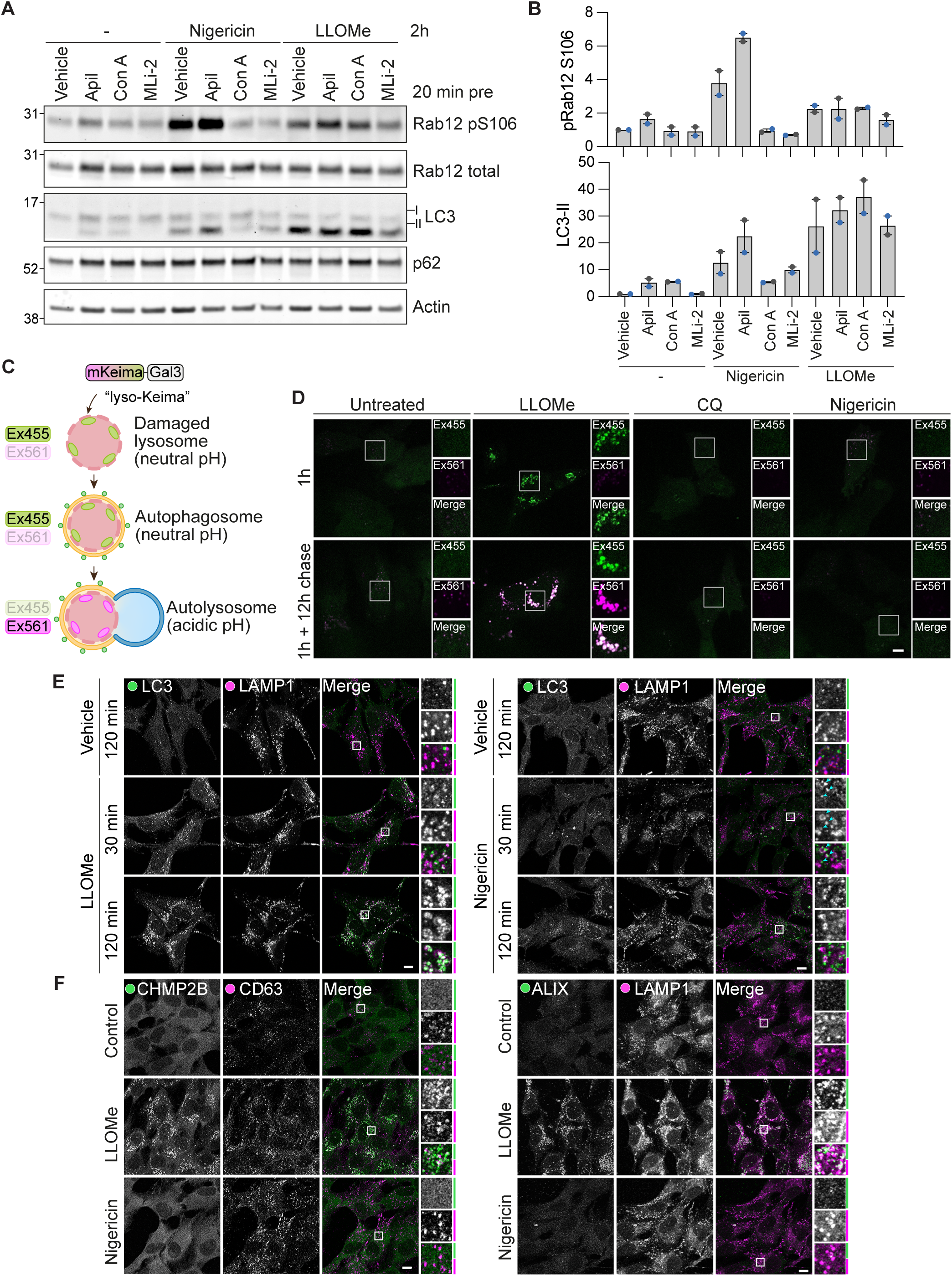
LLOMe, nigericin and chloroquine cause distinct responses to endolysosomal stress. **A.** Representative western blot of RPE1 Flp-In Parental cells pre-treated with Vehicle (DMSO), 150 nM apilimod (Apil), 100 nM concanamycin A (ConA) or 100 nm MLi-2 then treated with Vehicle (DMSO), 500 *µ*M LLOMe or 2 *µ*M nigericin for 2 hours prior to lysis. **B.** Quantification of A. Values normalised to vehicle control. n = 2. Error bars indicate mean and range. **C.** Schematic of mKeima-Gal3 (lyso-Keima) reporter principle **D.** Representative images of RPE1 Flp-In VPS35 WT cells (not induced with doxycycline) expressing mKeima-Gal3 treated for 1 h with LLOMe, nigericin or chloroquine (CQ) and then imaged immediately or imaged following a 12 h washout. Scale bar 10 *µ*m. **E.** Representative images of RPE1 Flp-In Parental cells treated with LLOMe or vehicle (DMSO) and nigericin or vehicle (EtOH) for the indicated times then fixed and stained for LC3 and LAMP1. Scale bar 10 *µ*m. **F.** Representative images of RPE1 Flp-In Parental cells treated with LLOMe or nigericin for 30 min then fixed and stained for markers for the ESCRT machinery (CHMP2B, ALIX) and late endosomes/lysosomes (CD63, LAMP1). Scale bar 10 *µ*m.

**Figure S2:**
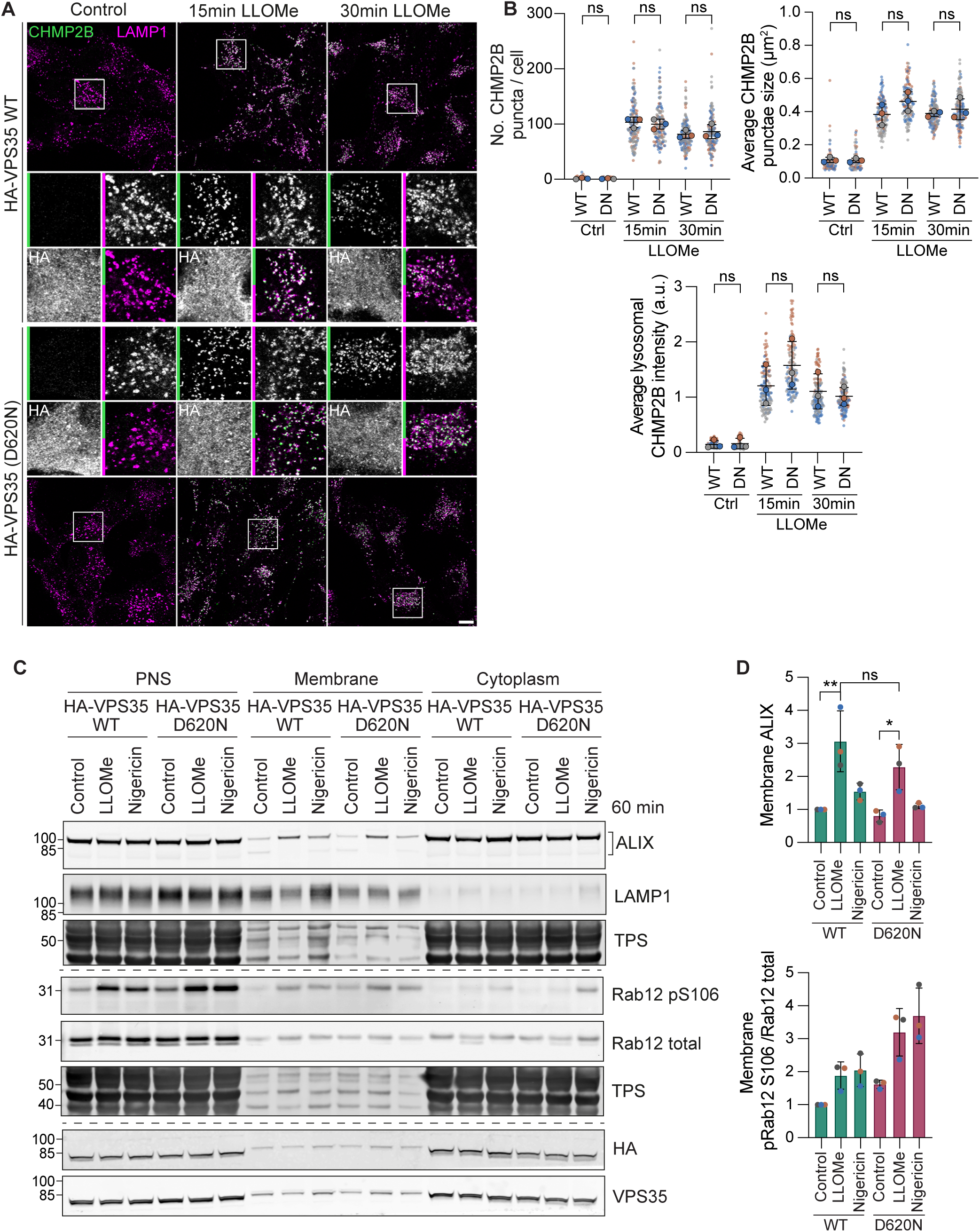
The VPS35 (D620N) mutation does not affect ESCRT recruitment in response to lysosomal membrane damage. **A.** Representative images of induced RPE1 Flp-In VPS35 WT and (D620N) cells treated with 500 *µ*M LLOMe for 15 or 30 min prior to fixation and staining with the indicated antibodies. Scale bar 10 *µ*m. **B.** Quantification of the number and average size of CHMP2B puncta per cell and the mean CHMP2B intensity within LAMP1 puncta per cell. 37-60 cells quantified per condition, n = 2. One-way ANOVA with Tukey’s multiple comparisons test. ns, not significant. **C.** Subcellular fractionation of RPE1 Flp-In VPS35 WT and D620N (DN) cells induced with 0.1 μg/ml doxycycline for 24 h and then treated with 500 *µ*M LLOMe, 3 *µ*M nigericin or vehicle control (DMSO) for 60 min. PNS; post nuclear supernatant, TPS; total protein stain **D.** Quantification of D. Values normalised to WT control. n = 3. Error bars indicate mean ± SD. One-way ANOVA with Tukey’s multiple comparisons test. P * < 0.05, P ** < 0.01

